# Accurate Detection of Incomplete Lineage Sorting via Supervised Machine Learning

**DOI:** 10.1101/2022.11.09.515828

**Authors:** Benjamin Rosenzweig, Andrew Kern, Matthew Hahn

## Abstract

Gene tree discordance due to incomplete lineage sorting or introgression has been described in numerous genomic datasets. Among distantly related taxa, however, it is difficult to differentiate these biological sources of discordance from discordance due to errors in gene tree reconstruction, even when supervised machine learning techniques are used to infer individual gene trees. Here, rather than applying machine learning to the problem of inferring single tree topologies, we develop a model to infer important properties of a particular internal branch of the species tree via genome-scale summary statistics extracted from individual alignments and inferred gene trees. We show that our model can effectively predict the presence/absence of discordance, estimate the probability of discordance, and infer the correct species tree topology in the presence of multiple, common sources of error. While gene tree topology counts are the most salient predictors of discordance at short time scales, other genomic features become relevant for distantly related species. We validate our approach through simulation, and apply it to data from the deepest splits among metazoans. Our results suggest that the base of Metazoa experienced significant gene tree discordance, implying that discordant traits among current taxa can be explained without invoking homoplasy. In addition, we find support for Porifera as the sister clade to the rest of Metazoa. Overall, these results demonstrate how machine learning can be used to answer important phylogenetic questions, while marginalizing over individual gene tree—and even species tree—topologies.

## Introduction

Topological discordance among gene trees is found in nearly all phylogenomic datasets. Biological causes of discordance include both incomplete lineage sorting (ILS; Figure 1) and introgression (Maddison 1997), while technical sources of discordance include substitution rate heterogeneity and model misspecification. Rampant discordance has been described in recent radiations, such as cichlids (Ronco et al., 2021), horses (Jónsson et al. 2014), *Drosophila* (Pollard et al. 2006), and tomatoes (Pease et al. 2016), as well as in ancient radiations such as birds (Jarvis et al. 2014) and land plants (Wickett et al. 2014). When discordance is due to biological causes, phylogenetics provides powerful tools for understanding genotype-phenotype associations (Pease et al. 2016; Smith et al. 2020), as well as for distinguishing traits that have evolved more than once (homoplasy) from traits that have evolved once on discordant gene trees (hemiplasy; (Avise and Robinson 2008)). Therefore, determining whether gene tree discordance is due to biological factors is a key step in understanding trait evolution.

**Figure 1:**
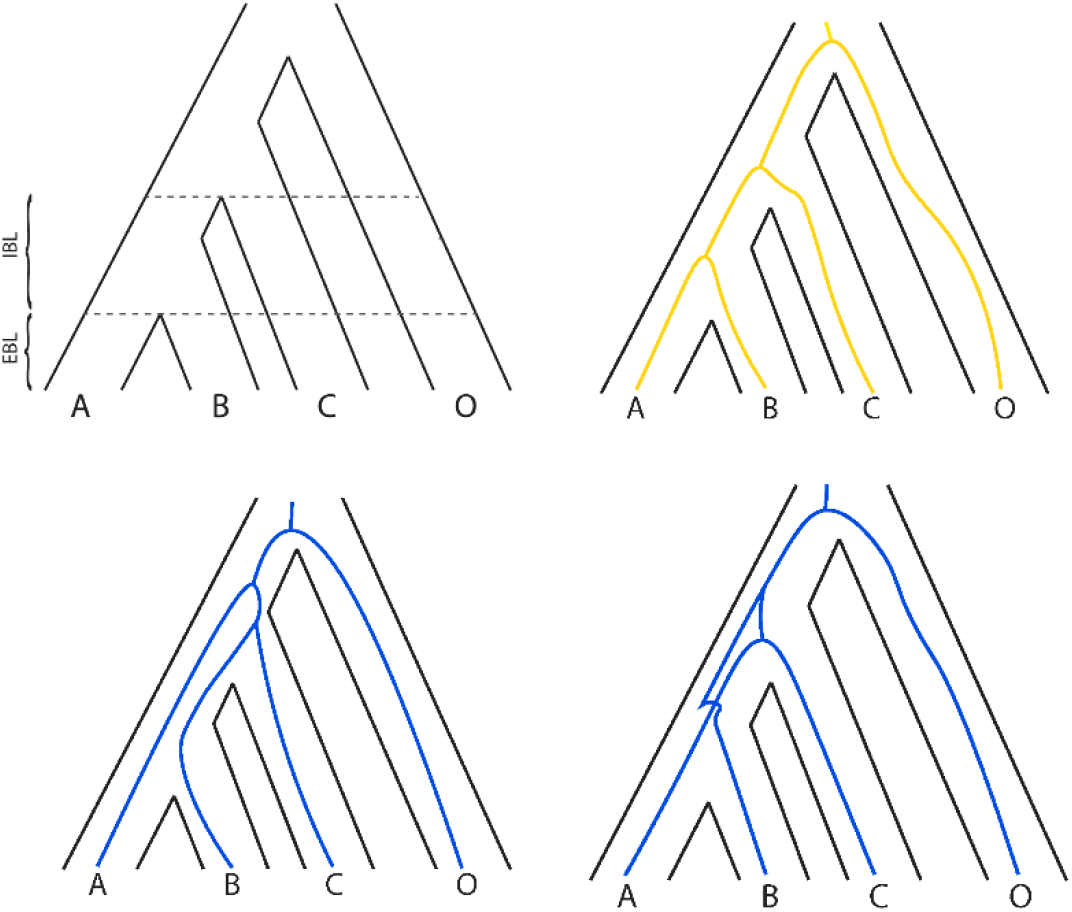
ILS among three taxa can produce one concordant (yellow) and two discordant (blue) gene tree topologies. The probability of concordance depends only on the internal branch length (IBL), while gene tree inference error depends on both IBL and external branch length (EBL).

For closely related species, individual gene trees can be recovered with high accuracy by maximum likelihood, Bayesian inference, or parsimony methods. For more distantly related taxa, gene tree reconstruction increasingly falls prey to errors due to homoplasy at individual sites (i.e. “multiple hits”). For truly ancient divergences, discordant topologies will be inferred at the same frequency as the species tree topology, regardless of the amount of discordance present due to biological factors (Figure 2). Evolutionary model misspecification, whether due to an incorrect substitution matrix or failure to account for site-and lineage-specific rate heterogeneities, compounds these errors. No method currently exists that can distinguish between true discordance—that due to biological processes—and the noise introduced by gene tree reconstruction error. Even when the species tree topology is known with high confidence, any assertion about the history of specific biological traits still requires accurately estimating the probability of hemiplasy, a procedure that requires knowledge of the level of true biological discordance among gene trees.

**Figure 2:**
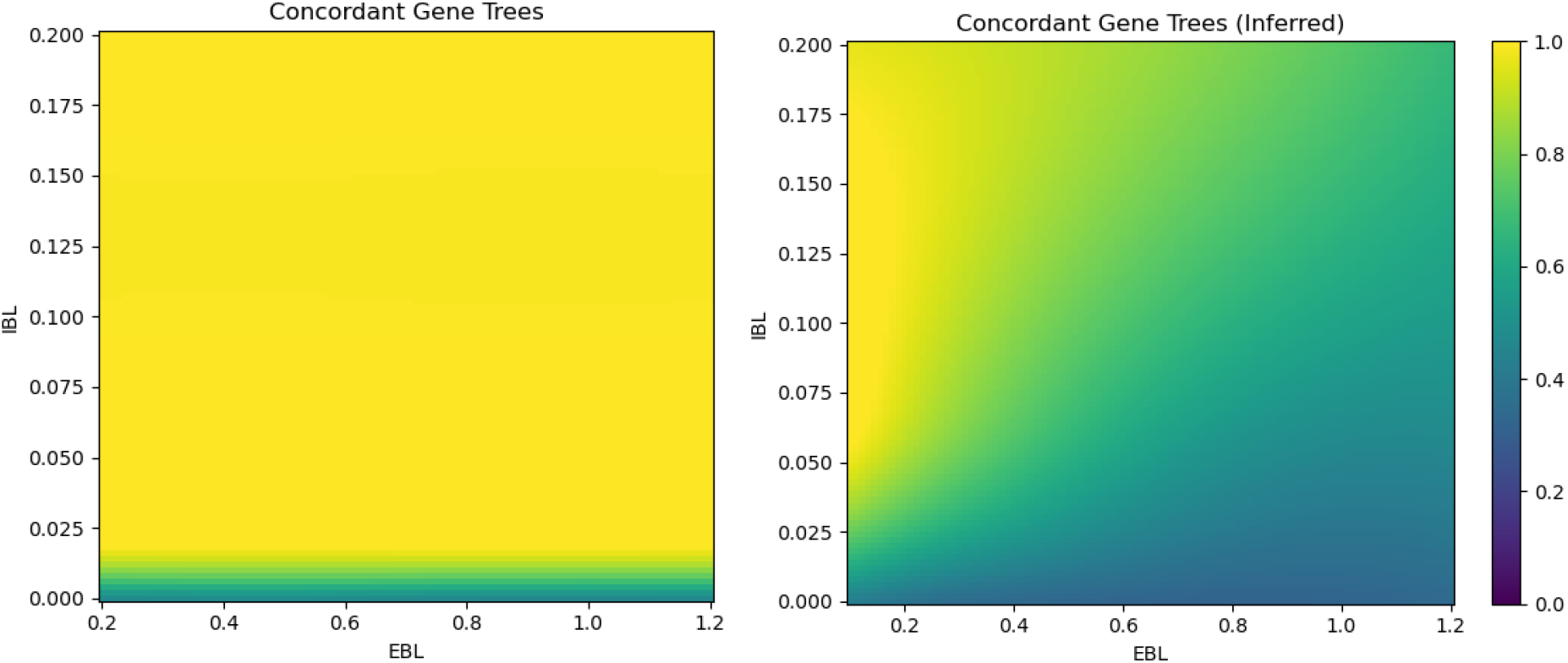
The fraction of true (left) versus inferred (right) gene tree topology frequencies that match the species tree topology (*p*) as a function of IBL and EBL. X- and Y-axes are in units of expected (amino acid) sequence divergence.

A particularly contentious problem in phylogenetics involves relationships at the base of the Metazoa. Over the last decade, genomic data has led systematists to reconsider the relationship between the ParaHoxozoa clade (comprising the phyla Bilateria, Cnidaria, and Placozoa) and the two other major clades of animals: the phyla Ctenophora and Porifera (Laumer et al. 2019; Simion et al. 2017; Whelan et al. 2015)). In the absence of biological discordance, the Ctenophora-sister hypothesis implies that major features such as nervous systems (Moroz et al. 2014, 201), basement membranes, and the through-gut either evolved multiple times or arose once in the ancestor of all three groups and then were lost in the Porifera. While much effort has been devoted to investigating the impact of fast-evolving lineages and the choice of substitution model on species tree reconstruction at the base of Metazoa (Li et al. 2021), scant attention has been paid to the potential role of gene tree discordance in reconciling the evolution of complex traits during this period (King and Rokas 2017). In the presence of biological discordance, even complex traits can arise once and appear in multiple lineages without needing to invoke homoplasy (Hahn and Nakhleh 2016; Guerrero and Hahn 2018; Hibbins, Gibson, and Hahn 2020).

Supervised machine learning (SML) comprises a variety of parameter-rich regression algorithms that excel at learning nonlinear mappings from noisy, feature-rich data. SML methods have been successfully employed in a variety of contexts in phylogenetics, from inferring quartet topologies for single loci (Suvorov, Hochuli, and Schrider 2020; Zou et al. 2019), to enhancing nucleotide substitution model selection (Abadi et al. 2020), to tree-search proposal distributions (Azouri et al. 2021). Rather than attempting to use SML to improve the accuracy of individual gene trees, our goal here is to predict the fraction of gene trees in a dataset that are biologically discordant. Our model infers properties of an internal branch of the species tree given a collection of summary statistics from a set of gene trees. Given the noisy nature of gene trees inferred from deep divergences (Figure 2), SML offers a potentially powerful method for overcoming inference problems in such datasets.

In this paper, we show that a variety of SML algorithms can effectively distinguish biological discordance from gene tree inference error across a wide range of parameter space. We demonstrate that simple feed-forward artificial neural network architectures are successful at (1) predicting the species tree topology, (2) predicting the fraction, *p*, of biological discordance in a set of gene trees, and (3) detecting the presence or absence of biological discordance in a given dataset. We show that biological discordance can be identified under a wide range of biologically relevant scenarios, even when the assumptions of the training regime are violated.

## Methods

### Supervised Machine Learning Models

The regression and classification models were implemented in scikit-learn (Buitinck et al. 2013), with the exception of the deep neural network (DNN) models, which were implemented in pytorch (Mazza and Pagani 2021). Hyperparameters for the DNN architectures and optimization algorithm were selected via a Bayesian optimization search over 2000 candidate configurations. Three major categories of SML model were employed: Linear (regularized linear and logistic regression), Ensemble (random forest, adaptive boosting, and gradient boosting with decision trees), and a DNN with rectified linear activations. The DNN estimator of *p* (the expected frequency of concordant trees), DNN-Prob, and the species tree topology predictor, DNN-Top, were trained with mean squared logarithmic error and cross-entropy losses, respectively. The DNN classifier, DNN-Class, was obtained by thresholding the DNN-Prob regressor.

### Simulation Conditions

All models were trained on the same training dataset, comprising 1.5 million synthetic sets of alignments, each of which included variable sequence lengths, site-specific evolutionary rates, and substitution models. Parameters for simulated training data are described in Table 1 and the overall process is shown in Figure 3.

**Table 1:**
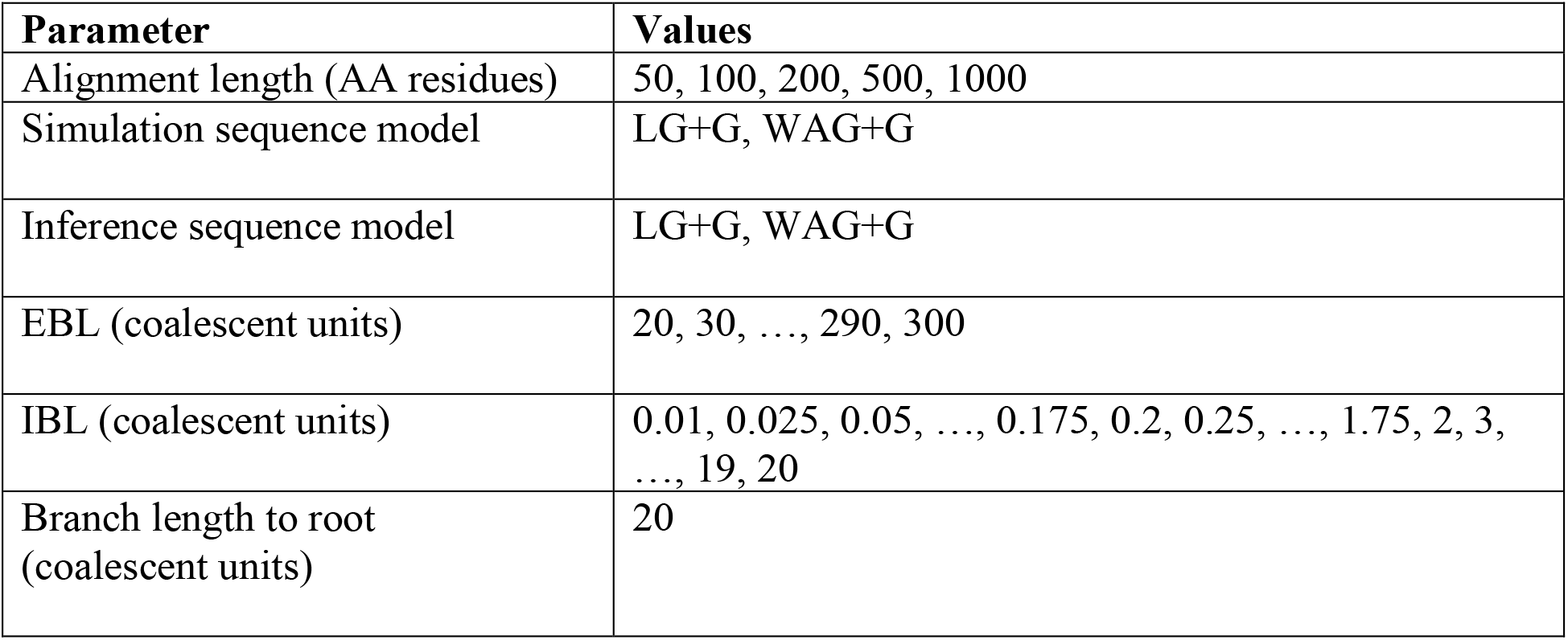
Simulated training conditions. At least 1,000 loci were simulated for each parameter combination.

**Figure 3:**
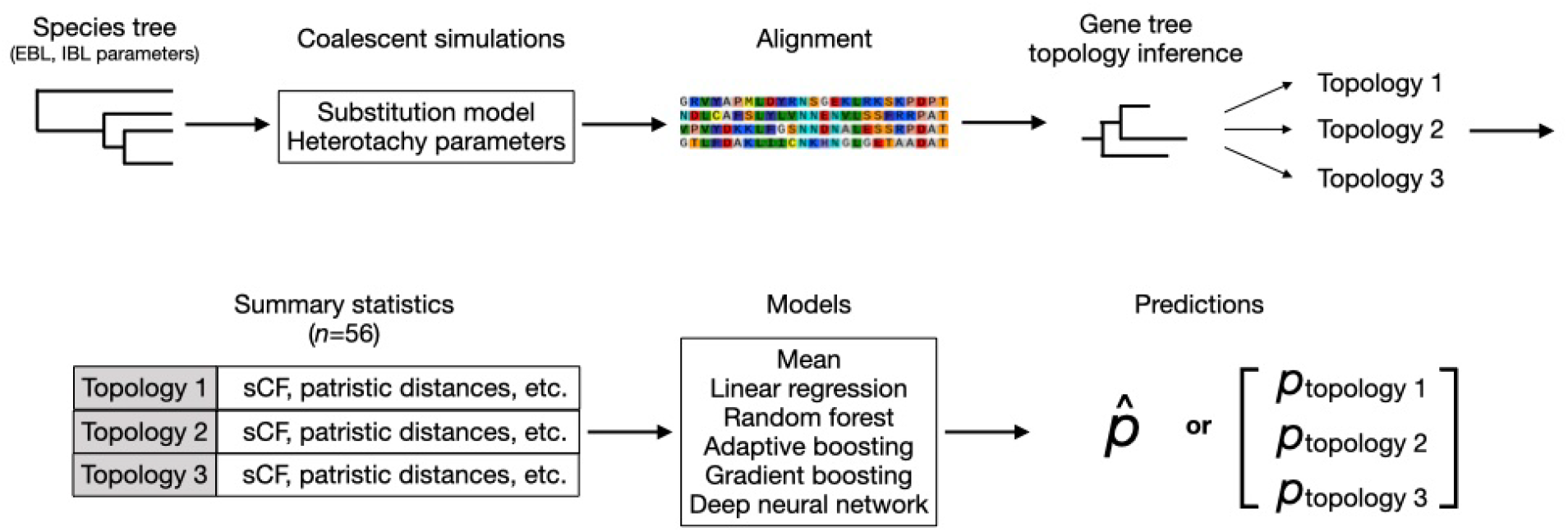
Flow chart of data processing pipeline. For a given set of species tree parameters, coalescent simulations are carried out in conjunction with the simulation of an associated alignment. For each alignment, gene trees are inferred, and sorted into bins by the inferred tree topology. For each of the 3 topology bins, 56 summary statistics are computed, for a final feature vector of length 168. A variety of supervised machine learning models are trained to predict either 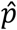, the probability of concordance (which can be used as a binary classifier with suitable threshold), or the probability of each species tree topology.

Four-taxon gene trees were simulated with ms (Hudson 2002) and alignments with seq-gen (Rambaut and Grassly 1997). For the training set, all simulated trees initially had the same rate of sequence evolution and all loci were simulated without recombination; below we describe how heterotachy and recombination were added in the test sets. Training sequences were simulated using both LG and WAG substitution models with a discrete 5-category gamma model of site-specific rate heterogeneity. Individual gene trees were inferred with RaXML-NG (Kozlov et al. 2019), using a discrete gamma model of site-specific rate heterogeneity with 4 categories and empirical amino acid frequencies. Site concordance factors were computed with IQ-TREE (Minh, Hahn, and Lanfear 2020).

For a given set of parameters, alignments were generated for a large number of loci (a minimum of 1,000 for each parameter set), and gene trees were inferred for each alignment. Raw features such as alignment lengths, site concordance factors, and patristic distances were computed from both the inferred trees and raw alignments. These feature values were then binned according to the rooted topology of the inferred tree. For every bin, summary statistics (central moments and order statistics) were computed across all values of each feature. This process resulted in 168 total features per simulated dataset (Figure 3). Features such as concordance factors, patristic distances, and topology counts are not invariant to relabeling of taxa; training data was therefore augmented with label permutations corresponding to the three rooted species trees. At test time, predictions (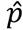 for the regression algorithms, softmax outputs for DNN-Top) were generated separately for each of the three label permutations of the feature vector, and the resulting predicted values were averaged across all three permutations (the input data were also permuted in a similar manner). The average of the ensemble was considered the point estimate of our model.

Additional conditions used for test datasets are described in Supplementary Table 1. Lineage-specific heterotachy was simulated by multiplying all ingroup external branches by independent Gamma(4, .25) multipliers. Recombination was simulated by randomly permuting subsequences within a synthetic dataset; this was done because simulations using the coalescent with recombination would have resulted in a training dataset that was too sparse. Both the number of recombination blocks per sequence and the fraction of loci experiencing recombination varied by dataset.

### Metazoa Datasets

We used 13 previously published datasets containing 315 taxa from three Metazoan clades (Supplementary Table 2). Most of these studies utilized concatenated alignments with either the general time reversible (GTR) substitution model across all sites or a form of data partitioning (Whelan and Halanych 2017). Our analysis requires individual gene trees, but individual genes do not provide enough data to reliably estimate GTR model parameters. We therefore used the LG substitution model (Le and Gascuel 2008) with Gamma-distributed site heterogeneity to infer gene trees for all data matrices. Our training set includes a mixture of substitution models (LG and WAG), as well as site heterogeneity. Although the training set did not include misspecified substitution models, the DNN methods are highly robust to substitution matrix misspecification (see next section). This justifies the use of a single substitution model across all gene trees for this analysis. Augmenting the training set with gene trees inferred from misspecified models could further improve predictive accuracy.

For each data matrix, we extracted alignments from all 4-taxon quartets comprising one member each from Ctenophora (Cte), Porifera (Por), ParaHoxozoa (PaH), and an outgroup from Choanoflagellata/Fungi/Ichthyospora/Filasterea (Out). For each gene, the corresponding subtree was rooted with the outgroup taxon. Quartets of taxa having fewer than 25 genes were excluded from the analysis. The distribution of inferred gene tree topologies from all quartets is displayed in Table 2.

**Table 2:** Number of quartets for which the most common gene tree topology supports Ctenophora-sister (C), Porifera-sister (P), or Parahoxozoa-sister (Pa). “Data Matrix” refers to the original papers from which datasets were obtained. For a full list and description of all datasets see Supplementary Table 2.

| **Data Matrix** | **C** | **P** | **Pa** |
| --- | --- | --- | --- |
| Borowiec2015_Best108 | 6 | **114** | 60 |
| Borowiec2015_Total1080 | 6 | **122** | 64 |
| Nosenko2013_ribo_11057 | 239 | **338** | 242 |
| Ryan2013_est | 203 | **854** | 470 |
| Ryan2013_est_only_choanozoa | 87 | **229** | 143 |
| Ryan2013_est_only_holozoa | 134 | **333** | 217 |
| Whelan2015_D10 | 15879 | **36816** | 23400 |
| Whelan2015_D10_only_choanozoa | 1916 | **8296** | 4647 |
| Whelan2015_D1_only_choanozoa | 1990 | **8488** | 4786 |
| Whelan2015_D1_only_holozoa | 6125 | **12223** | 7872 |
| Whelan2015_D20_only_choanozoa | 478 | **978** | 602 |
| Whelan2017_full | 250 | **542** | 342 |
| Whelan2017_full_only_choanozoa | 747 | **1068** | 747 |

## Results

### Predicting the probability of concordance

#### Predicting *p*

In our first set of experiments, we trained a variety of SML methods to predict *p*, the probability that a 4-taxon gene tree matches the topology of its species tree. The value of *p* ranges from 1/3 (all topologies are equiprobable) to 1 (all gene trees are concordant) and is a function of internal branch lengths (IBL; Hudson 1983). The quantity 1-*p* therefore represents the true probability of discordance in a dataset.

Figure 4 shows the error, 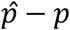, of all six SML models across a range of external and internal branch lengths (EBL and IBL). SML models differ in their error rates (Kruskal-Wallis test, H = 2747.88, p < 10^−300^); one-sided Wilcoxon rank-sum tests were conducted between each pair of algorithms to explore differences in performance. Gradient Boosting (GradBoost) and the deep neural network (DNN-Pred) performed best, followed by Random Forest (RF), then Adaptive Boosting (AdaBoost), then the constant baseline predictor (Mean; equal to 0.999995, the mean *p* across the training set), and finally the regularized linear regression (ElasticNet) (all *P* < 10^−80^, Holms-Bonferroni corrected α= 3.34 × 10^−4^). Although the overall error rate of DNN-Pred was not significantly better than GradBoost, its performance was more consistent: the 50% interquartile ranges of error for all algorithms were: 0.4043 (ElasticNet), 0.399 (RF), 0.4040 (AdaBoost), 0.0097 (GradBoost), 0.0006 (DNN-Pred).

**Figure 4:**
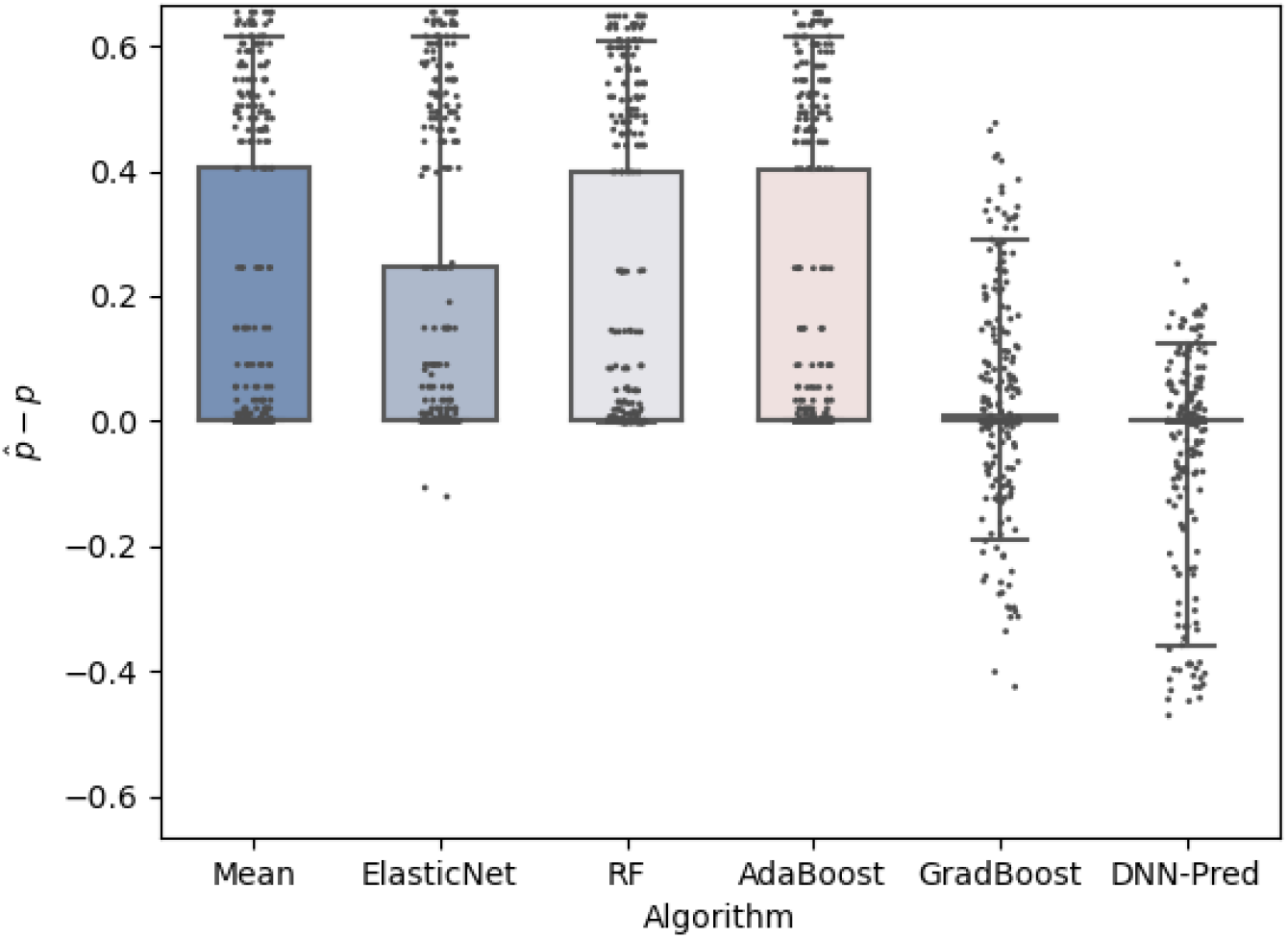
Error of all SML methods on a collection of 250-gene, 500-residue datasets. EBL ranges from 0.2 to 1.0, IBL from 0.001to 0.18. The baseline Mean predictor is the mean *p* across all samples in the training sets. The other predictors from left to right are: regularized linear regression (ElasticNet), random forest (RF), adaptive boosting with decision trees (AdaBoost), Gradient Boosting with decision trees (GradBoost), and the deep neural network predictor (DNN-Pred). Boxes represent the 25% and 75%, quantiles, whiskers the 5% and 95% quantiles.

In the remainder of the paper, we therefore focus on results from the deep neural networks. Figure 5 shows the performance of DNN-Pred on simulated test data as a function of EBL and IBL. 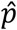 is most accurate when (a) both IBL and EBL are short or (b) IBL is long. For long EBL, 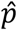 tends to be an overestimate of *p* for short IBL (where expected amino acid sequence divergence < 0.01) and an underestimate for intermediate IBL (between .01 and 0.05). As expected, performance is best when IBL is extremely long (> 0.07, corresponding to *p* > 0.99999). In this region gene tree inference error is negligible even for long EBL. Underestimating *p* for IBL in the 0.01 to 0.05 range is a feature shared with the gradient boosting algorithm, the only other method with reasonable performance across the regions of parameter space we explored.

**Figure 5:**
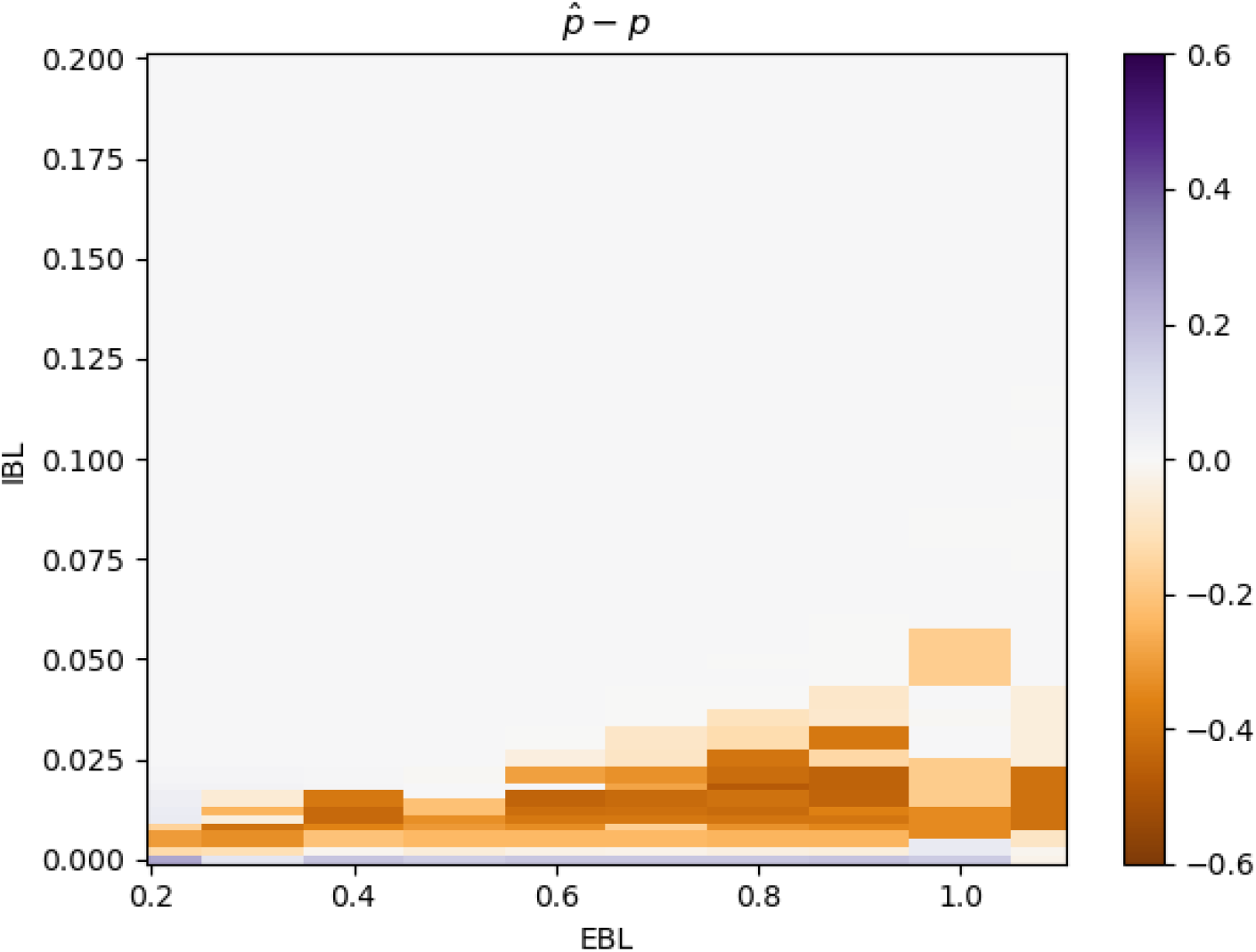
Accuracy of DNN-Pred. Average distance between true and predicted *p* for 500-gene, 500-residue datasets. Branch length units on both axes are in expected amino acid sequence divergence.

### Model Misspecification

#### Recombination

Long protein-coding genes have almost certainly experienced recombination, and therefore combine multiple topologies (Lanier and Knowles 2012). This increases the risk of hemiplasy (Mendes, Livera, and Hahn 2019), leading to inaccurate estimates of both divergence times and tree topologies. To investigate the impact of recombination on prediction accuracy we performed additional manipulations. To mimic the presence of intra-locus recombination, we varied the fraction of recombinant loci present in each dataset (Figure 6), drawing each recombinant locus uniformly from 2, 3, and 4 independently sampled blocks of sequence.

**Figure 6:**
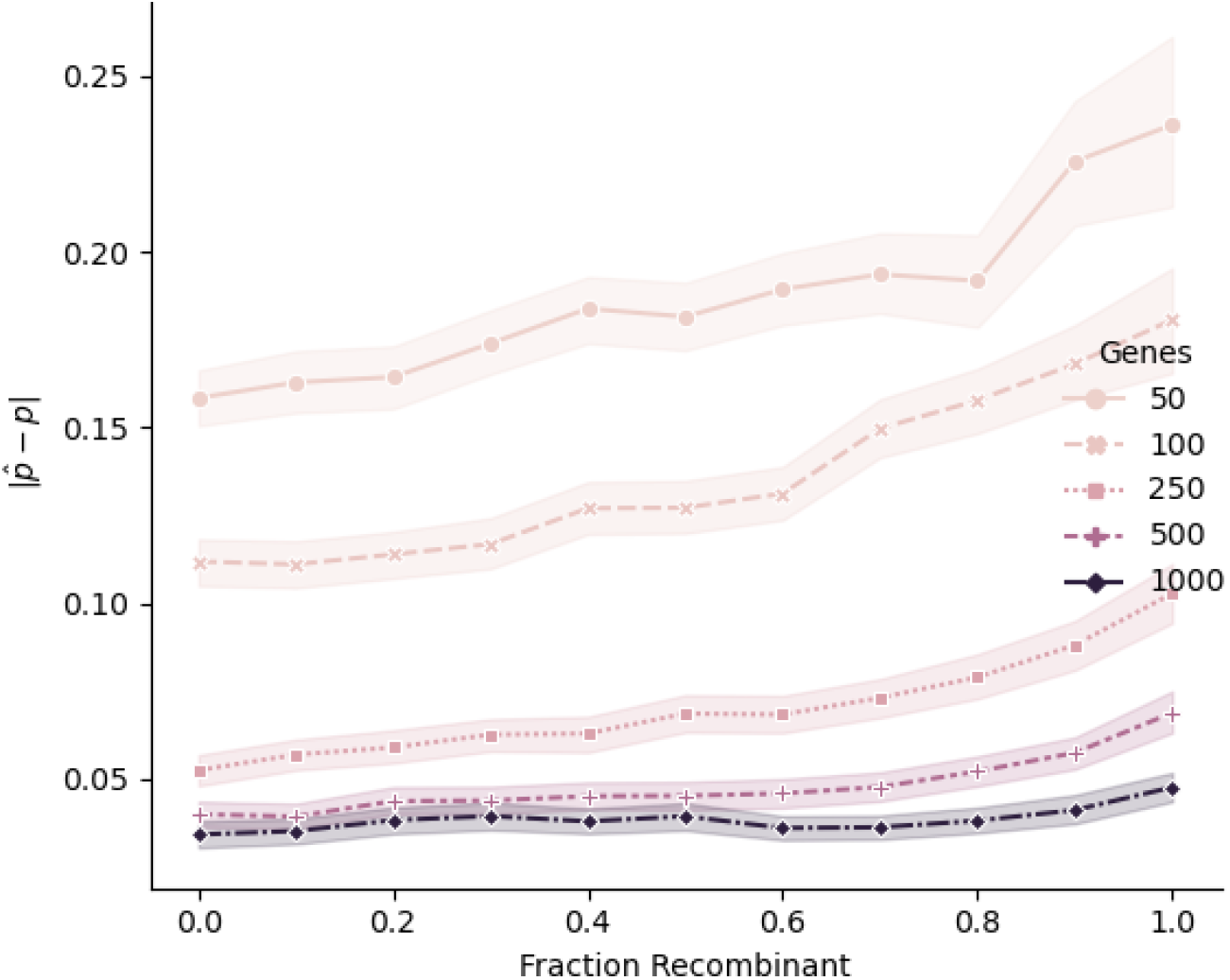
Intragenic recombination reduces the accuracy of *p*. Average absolute error between true and predicted *p* for datasets of length 500 residues is shown across a range of parameter space. Recombinant genes are drawn uniformly from 2-, 3-, and 4-block conditions, and both the number of genes and fraction of recombinant genes per test set are varied.

Although increased intragenic recombination has an effect on DNN-Pred’s ability to predict *p*, dataset size can effectively overcome this effect. For any single dataset size, we do see increasing error with an increasing fraction of recombinant sequences (Figure 6), but also increased error for smaller datasets even in the absence of recombination. To understand why this is the case, note that DNN-Pred relies heavily on site concordance factors and gene tree topology counts for much of parameter space (see Section “Feature Importance”). These features are minimally impacted by intragenic recombination: site concordance factor-derived statistics are pooled across genes and, while recombination may cause branch lengths of individual gene trees to be overestimated (Mendes & Hahn, 2016), the topology is still more likely to match the “majority tree” of the constituent sequences, than either minor tree.

#### Evolutionary model misspecification

We investigated the impact of several forms of evolutionary model misspecification: differing substitution matrices in training and test data, and hidden site- and lineage-specific rate heterogeneity. Each of these cases can be thought of an instance of what is called “data drift” in the machine learning world, where training data does not match empirical data which we wish to perform inference on. Misspecification of the substitution matrix has a negligible effect on predictive accuracy: prediction error on datasets simulated with the LG substitution matrix and inferred with WAG, or vice versa, do not differ significantly compared to datasets where trees were inferred with the correct substitution matrix (Supplementary Figure 1a). This suggests that sequence-based features (i.e. site concordance factors) provide enough information to estimate *p* despite errors in gene tree inference, or that substitution model misspecification does not systematically bias gene tree inference error in a way that obscures information regarding *p*. Further, to deal with rate heterogeneity we trained using simulations that included variation in rates among sites. This allowed for successful inference in the face of both site-specific heterogeneity and lineage-specific heterogeneity (heterotachy) in test data and did not significantly increase the error in estimating *p* (Kruskal-Wallis test; *H* = 0.686, *P* = 0.710; Supplementary Figure 1b).

### Binary Classification and Species Tree Prediction

In regions of parameter space where *p* cannot be inferred precisely, it is still useful to know whether *any* ILS exists in a particular dataset. Statistical power can be increased by introducing a binary classifier (DNN-Class) which predicts whether *p* is above or below a particular value. Note that distinguishing between the events *p* < 1 (“potential ILS”) and *p* = 1 (“no ILS whatsoever”) is not informative, as *p* = 1 corresponds to an infinitely long internal branch. We instead chose a target value of 0.9: this is interpreted to mean that datasets with *p* < 0.9 exhibit biological discordance and those with *p* > 0.9 do not. A more (or less) conservative target value than 0.9 can be chosen depending on the dataset. (For a dataset of *N* genes, for example, choosing a target value of 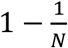 would ensure that the expected number of discordant trees under the “no ILS” hypothesis will be less than 1). DNN-Class was constructed by applying a separate decision threshold to the output of DNN-Pred. The decision threshold of 0.9511 was chosen to optimize the geometric mean of the classifier’s precision and recall on the training set (Figure 7a); a different target value will result in a different decision threshold.

**Figure 7:**
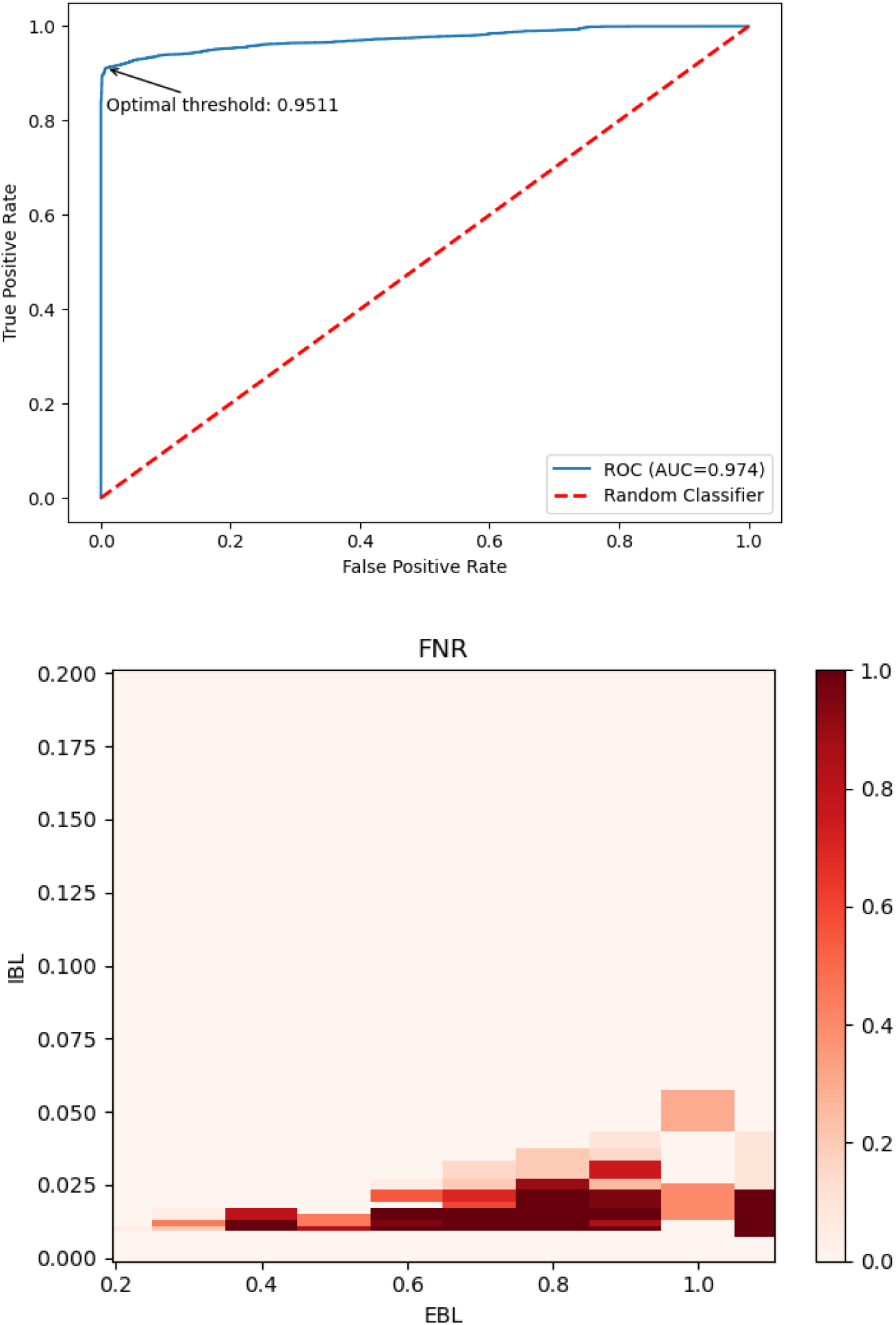
Performance of DNN-Class on 500-gene, 500-residue datasets. a) Receiver Operating Characteristic (ROC) curve. The area under the curve (AUC) is also reported. b) False negative rate (FNR) at the optimal threshold in panel a) as a function of internal and external branch length. Branch length units on both axes are in expected amino acid sequence divergence.

Although a task-specific DNN classifier can be trained independently (e.g. with a binary cross-entropy loss), we found that this did not improve over the procedure described above. At this threshold, the probability of incorrectly predicting no ILS (“false positives”) is negligible for all parameters tested. The probability of incorrectly predicting ILS (“false negatives”) is below 0.3 in most regions of parameter space, with an elevated false negative rate for extreme values of EBL (Figure 7b). While DNN-Pred tends to overestimate *p* for IBL<.01, DNN-Class achieves zero error in this region. These results support the use of DNN-Class as a conservative estimator of concordance (i.e. “no ILS”).

Similarly, we can construct a ternary classifier (DNN-Top) to predict which of the three possible topologies is most likely to be the underlying species tree. The accuracy of DNN-Top varies with EBL and IBL in a pattern similar to DNN-Pred and DNN-Class (Figure 8). When IBL is greater than 0.0015 (corresponding to *p* ≈ 0.506), DNN-Top can correctly infer the species tree topology with greater than chance accuracy for all values of EBL tested. For IBL greater than 0.025 (*p* ≈ 0.996), accuracy is greater than 90%, even when a large fraction of gene trees are incorrectly inferred due to inference error.

**Figure 8:**
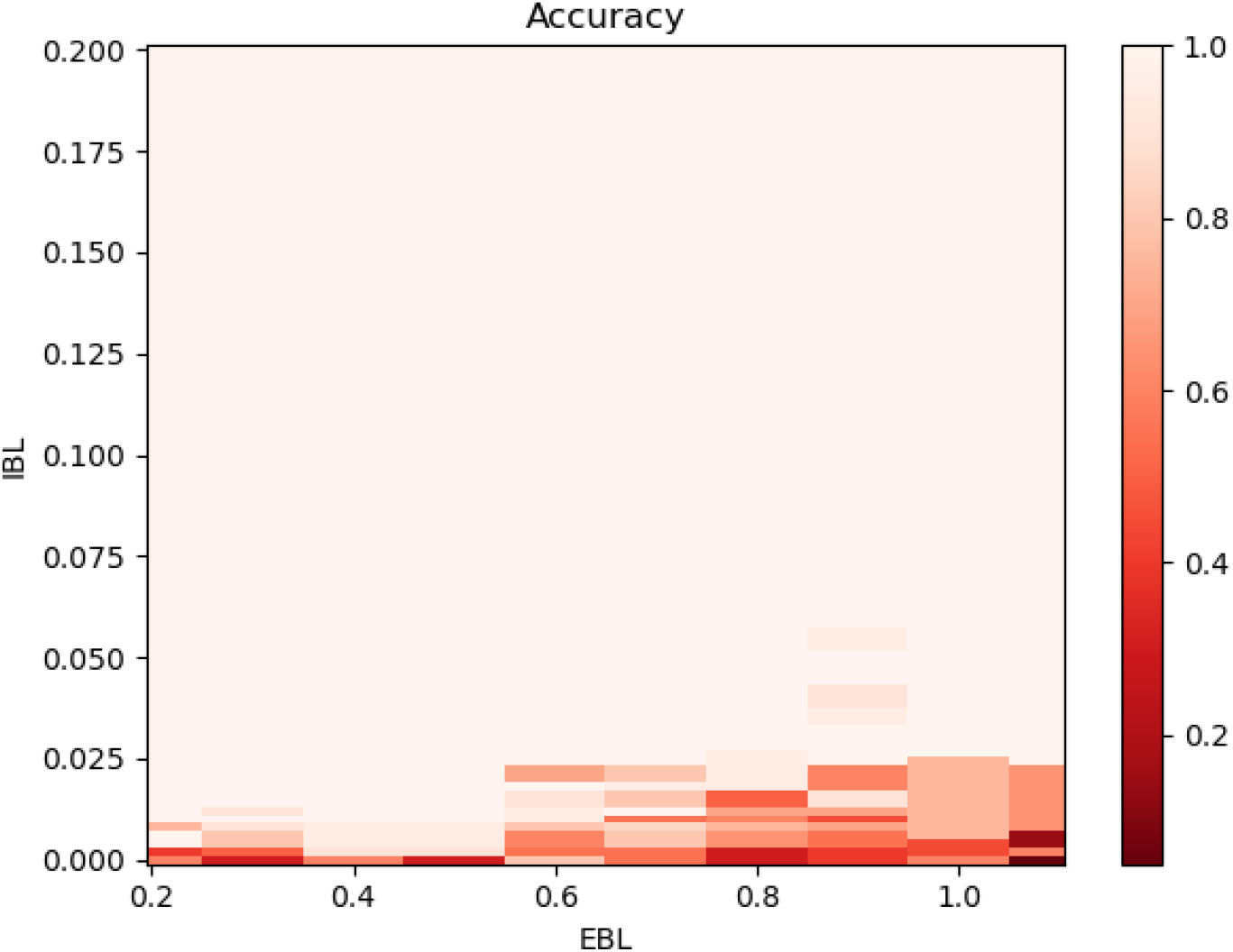
Accuracy of DNN-Top on 500-gene, 500-residue datasets. Accuracy is measured as the fraction of topology predictions that match the true species tree topology. Branch length units on both axes are in expected amino acid sequence divergence.

### Feature Importance

Tree-based classification and regression algorithms enable an intuitive notion of feature importance: the extent to which each covariate contributes to a prediction can be measured by the *Gini importance*: the mean decrease in impurity (for classification trees) or variance (for regression trees) at each node in which that covariate appears (Louppe et al. 2013).

Supplementary Figure 2a shows the features with the highest Gini importance for the gradient boosting regressor. Topology counts and sCF-derived features are the most informative features. This is expected, as topology counts alone are sufficient to recover *p* in the short external branch regime.

Untangling the importance of features in a DNN is considerably more challenging, and this opacity has hampered the entry of DNN’s into many fields of research. One method that has recently gained in popularity is the Shapley value: the average marginal contribution of each feature across all subsets of features. Exact computation of Shapley values is linear in the number of instances and exponential in the number of features, making it impractical for large, high-dimensional datasets. Numerous approximate methods have therefore been developed. We utilized Deep SHAP (Lundberg 2017), an approximate Shapley algorithm that leverages the compositional nature of neural networks. Like other measures of feature importance, the Shapley method assumes features are uncorrelated. This is not the case in our model, both because certain summary statistics for each feature are highly correlated (all measures of dispersion or of central tendency), and because the symmetry of the data makes certain combinations of features equivalent (permutations of taxa labels should not affect the output of DNN-Pred or DNN-Class). To overcome this, we computed Shapley values for the following groups of features: statistics derived from pairwise patristic distances *d(i,j)* (15 features each), statistics derived from counts of informative sites (15 features each), statistics derived from alignment lengths (15 features each), statistics derived from site concordance factors (45 features) and counts of gene tree topologies (3 features). As with tree-based regressors, topology counts and site concordance factors are the most informative features at short distances (Supplementary Figure 2b). For deeper divergences, features derived from gene tree branch lengths become more important (Supplementary Figure 2c). Unlike the gradient boosting regressor, DNN-Pred relies heavily on sequence length and “number of informative sites” statistics. As these variables determine the quality of other features (such as the inferred gene tree topologies and patristic distances), they may be important in determining how the network weights such features for a given instance.

### Biological Discordance at the Base of the Metazoa

In order to highlight the power of the SML approach described here, we applied our DNN models to a dataset in which the presence of discordance would have important biological implications. We used 13 datasets containing 315 taxa from three Metazoan clades to examine discordance at the base of animals. Because these datasets contain many more than four taxa, for each dataset we analyzed many possible sampled quartets: one species from each of Ctenophore, Porifera, ParaHoxozoa, and an Outgroup, using all genes shared by these four sampled species. Each quartet was analyzed using DNN-Pred (“how much discordance is there?”), DNN-Class (“is there *any* discordance?”), and DNN-Top (“what is the species tree?”).

#### Level of discordance

Using DNN-Pred, estimates of *p* were highly variable across quartets both between datasets and within each dataset (Supplementary Figure 6). The distribution of 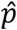 values was bimodal, with peaks at 0.43 and 0.88); this variability may be due to underlying variability in the rate of evolution of individual genes or taxa sampled for each quartet. Taking 0.6 as the putative metazoan EBL (one half the average patristic distance between the two sister taxa across all studies), the prediction error of DNN-Pred on simulated data in this area of parameter space ranges from -0.022 to 0.17 as a function of the true (unknown) IBL (Figure 5); this corresponds to a relative error of -4% to +50%, at the base of metazoa. Given this level of prediction error and putative values of 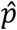, we cannot be certain using DNN-Pred whether there is truly biological discordance in this dataset. In trees with more recent common ancestors, 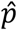 will be much more accurate (Figure 5).

#### Presence of discordance

In contrast, at this same level of divergence, the maximum false negative rate and false positive rate of DNN-Class are 0.20 and 0.00, respectively, across all IBL. This gives us confidence that the presence or absence of ILS can still be detected even if the expected frequency of concordant gene trees cannot be precisely determined. In all but one dataset, DNN-Class infers *p* < 0.9 in the overwhelmingly majority of quartets, providing strong evidence for ILS at the base of Metazoa (Figure 10a). The results from DNN-Class suggest that significant amounts of discordance are due to ILS, though we cannot say precisely how much.

**Figure 10:**
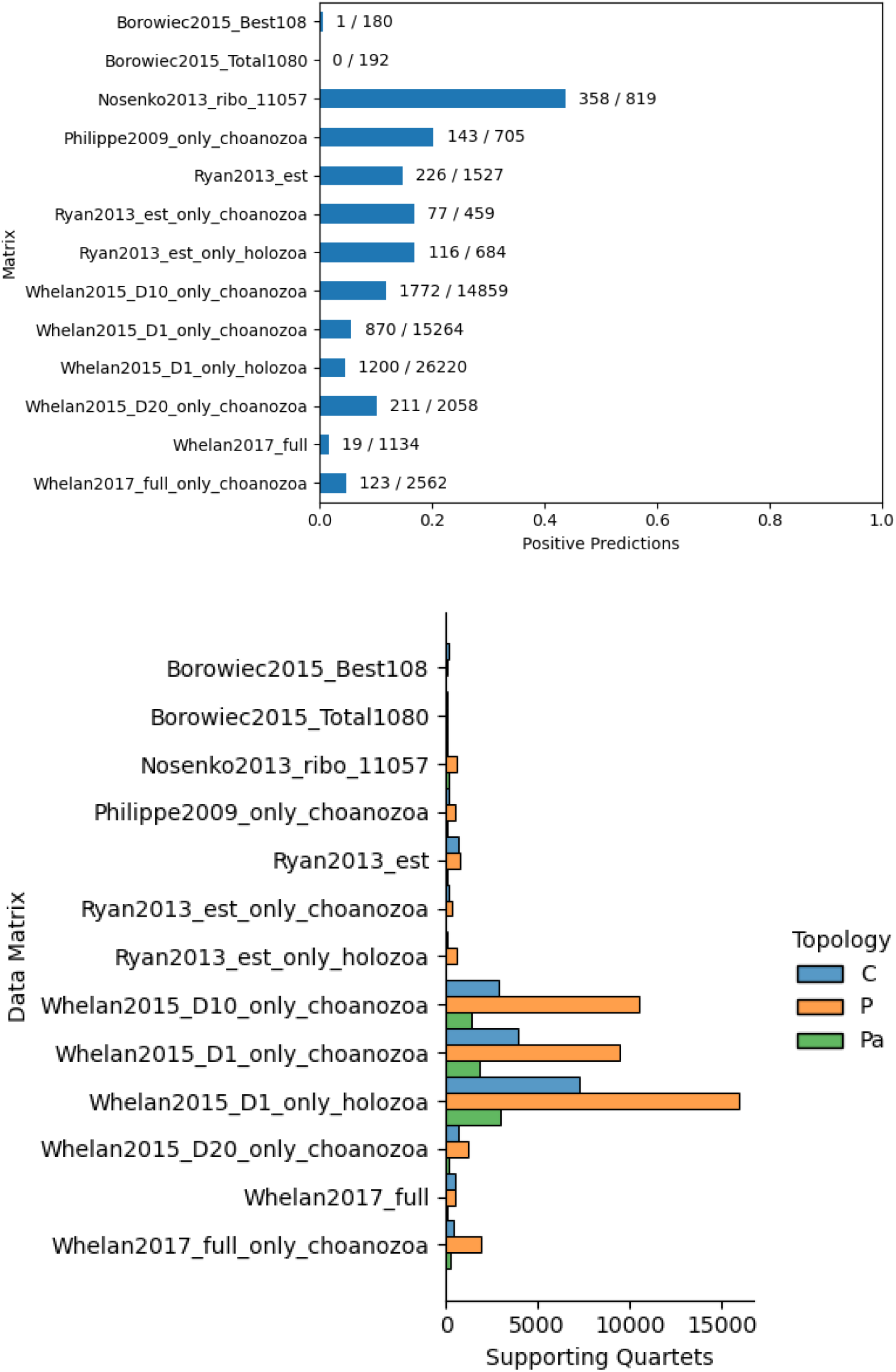
Discordance in the Metazoa. a) The fraction of cases with no discordance (DNN-Class predicts *p* > 0.9) across all Cte/Por/Par/Out quartets for each of 13 Metazoan data matrices. The number of such cases and total number of quartets in each dataset is also shown. b) Species tree topologies predicted by DNN-Top for the Metazoa datasets. Across all data matrices, the majority of quartets support the Porifera-sister species topology.

Looking within datasets, quartets in which DNN-Class predicted low *p* (high probability of ILS) have significantly more informative sites (two-sided *t*-tests, *P* < 10^−8^) and have higher site concordance factors (sCFs) for alternate topologies (two-sided *t*-tests, *P* < 10^−4^), assuming that the species tree is Porifera-sister (see next section). Sequences with more informative sites carry more phylogenetic signal; feature values and predictions for these quartets should therefore be considered more reliable. The only dataset with a significant number of quartets supporting *p* > 0.9 is the Nosenko et al. (2013) ribosomal dataset. With the exception of some sCF statistics, all feature values for this dataset differed significantly from the other datasets examined (Mann-Whitney U tests with Holm-Sidak correction, all *P* < 0.001). The features which varied most from other datasets were those derived from sequence length and pairwise distances involving the outgroup; the Nosenko et al. dataset has much shorter sequences than the other datasets examined, as well as longer outgroup branch length statistics (with sequence lengths at least 1.5 standard deviations below and branch lengths at east 1.5 standard deviations above the global mean). This suggests that stochastic error (due to short sequences) and systematic error (due to long branch attraction) may be especially high in this dataset, making these predictions less reliable.

#### Species-tree prediction

Across all datasets, there was consistent support for the Porifera-sister topology using predictions from DNN-Top (Figure 10b; Table 2). The output of DNN-Top is in the form of posterior probabilities for each topology, and therefore model confidence can be assessed by comparing such values. Site concordance factors are strongly associated with model confidence, measured as the absolute value of the log-likelihood ratio of the Porifera-sister and Ctenophora-sister hypotheses: quartets for which DNN-Top strongly favors Porifera-sister have high sCF values for the Porifera-sister topology; likewise quartets for which DNN-Top favors Ctenophora-sister have high sCF values for Ctenophore sister (Supplementary Figure 8).

## Discussion

Topological discordance is a ubiquitous feature of phylogenomic datasets. When gene trees are accurately inferred, this discordance can be attributed to biological factors such as ILS or introgression. In this study we find that SML algorithms can provide useful information regarding ILS even in the presence of gene tree reconstruction error. Our estimators perform well even in worst-case scenarios, where test data contain multiple sources of noise not present in the training data. Improvements in specific cases can be obtained by retraining models with simulated data that approximates the suspected sources of noise in a target dataset, for instance in the manner that we trained under site-specific rate heterogeneity.

Previous applications of SML in phylogenetics have reported results from a single algorithm or neural network architecture, presumably the result of a laborious trial-and-error process that often goes unreported. Here we take a systematic approach to evaluating a range of SML algorithms, tuning their hyperparameters, and characterizing their performance and limitations. Given the well-justified opposition to adopting “black box” inference models in systematics (and biology more generally), this analysis provides a useful framework for future SML studies to emulate and improve upon. Recent studies have used convolutional neural networks to infer 4-taxa topologies for individual gene trees from multi-sequence alignments (Suvorov, Hochuli, and Schrider 2020; Zou et al. 2019; Solis-Lemus, Yang, and Zepeda-Nunez 2022). Although these approaches compare favorably with conventional approaches to topology inference, all methods fail on deep divergences, presumably due to the same recurrent mutation phenomena that confound likelihood, parsimony, and distance-based methods (Molloy and Warnow 2018). Furthermore, extending such methods to phylogenies with more than four taxa requires some form of tree reconciliation (Strimmer and von Haeseler 1996; Snir and Rao 2012), which has been shown to be less accurate than standard Neighbor-Joining or maximum likelihood inference (Zaharias, Grosshauser, and Warnow 2022). Here, we take a different approach: we develop SML methods to infer important properties of a particular internal branch in the species tree via genome-scale summary statistics extracted from individual alignments and inferred gene trees. This approach allows us to utilize state-of-the-art maximum likelihood methods to infer gene trees from large multisequence alignments, which can then be subsampled to obtain quartets that contain the branch of interest. This method provides enough signal to resolve a problematic internal branch and to accurately classify a dataset as containing biological discordance, even when inferred gene trees contain large amounts of noise.

The approach taken here also demonstrates the advantage that SML can provide over traditional methods in answering highly tailored questions in phylogenetics. To this end, we emphasize that *p* is a property of the dataset as a whole—our methods do not predict whether an individual gene tree is concordant or discordant. Future research could likewise focus on training SML models to predict specific quantities of interest from raw data, rather than separately optimizing each step of an analysis pipeline or trying to solve every tree inference problem with a single model. SML with deep learning has been successfully applied in other phylogenomic studies to distinguish between reticulated and bifurcating phylogenies (Burbrink and Gehara 2018), between different hybridization scenarios (Blischak, Barker, and Gutenkunst 2021), and to identify selective sweeps in population genomic datasets (Isildak, Stella, and Fumagalli 2021; Kern and Schrider 2018). A major disadvantage of all such approaches (including our own) is the large amount of training data required. When this data is produced via coalescent simulations as in the present study, this translates into months of compute time (∼65 CPU-hours to generate 1000 gene trees, simulate their alignments, and infer maximum likelihood trees on a 2.5 GHz processor). Furthermore, our results show that a great deal of simulation and training resources are expended in exploring insoluble regions of parameter space: those areas where even the richest models with the largest training samples can do no better than a random classifier. Future SML applications might consider utilizing some form of active learning (Settles 2012) to simulate and train jointly, focusing effort on those regions where learning is tractable and ignoring regions where SML does not appear to provide any advantage.

Summary methods for species tree inference such as ASTRAL (Zhang et al. 2018), MP-EST (Liu, Yu, and Edwards 2010), and ASTRID (Vachaspati and Warnow 2015) are statistically consistent when given accurately inferred input gene trees, even in the presence of ILS. These methods fail, however, when phylogenetic signal is obscured by high levels of gene tree reconstruction error (Molloy and Warnow 2018). The fact that DNN-Top and DNN-Class perform well across a similar range of parameter space suggests that that deep neural networks may be capable of inferring species trees along with their internal branch lengths in a manner similar to summary methods, and may even be able to improve upon them by being more robust to gene tree error. Future work could focus on modeling this gene tree error before it is input into summary methods.

Applying our method to relationships at the base of the Metazoa, we find strong support for both the Porifera-sister hypothesis and for the presence of appreciable amounts of incomplete lineage sorting. Branch-specific heterotachy has been cited as a key factor in the uncertainty regarding the base of Metazoa (Pisani et al. 2015; Kapli, Yang, and Telford 2020). In particular, the long branch leading to Ctenophora has prompted many researchers to explore amino acid recoding strategies to uncover hidden phylogenetic signal (Redmond and McLysaght 2021). While there are several important caveats to the inferences presented here, we have shown that our methods (i.e. DNN-Top) are robust to heterotachy, as well as a number of additional model violations. We therefore conclude that there is good reason to believe that we are accurately classifying the species topology.

Even if a species tree topology is known with high confidence, any assertion about the history of specific biological traits still requires accurately estimating the probability of hemiplasy – a procedure that requires knowledge of the level of true biological discordance among gene trees. Much of the argument around the “true” set of relationships at the base of the Metazoa has presented alternative species topologies as uniquely tied to alternative trait histories. Despite our apparent inability to accurately estimate the exact level of biological discordance this deep in time (i.e. DNN-Pred), even the consideration of hemiplasy means that the species topology can be separated from trait histories. In this sense, the results from DNN-Class should play an important role in understanding the complex pattern of events that occurred during the early stages of animal evolution. Importantly, our results concluding that there was biological discordance during this period do not rely on any particular species tree topology, nor do they necessarily imply that discordance is due to ILS. Although it would be rash to make definitive conclusions at this time, the asymmetry in frequency of the two minor gene tree topologies (Table 2) and predicted species tree topologies (Figure 10b) could be due to introgression among early animals (cf. Huson and Bryant 2006). Given the ubiquity of introgression across the tree of life, there is no reason to think it was not also occurring early in animal evolution, possibly affecting key biological innovations.

## 5 Acknowledgements

This research was supported by a Department of Defense SMART fellowship (BKR) and National Science Foundation grant DEB-1936187 (MWH). ADK was supported under NIH award 1R01HG010774.

## 6 Supplementary Material

Code used in this paper is available at https://github.com/bkrosenz/ml4ils.

