## Supplementary Figures & Tables for "Accurate Detection of Incomplete Lineage Sorting via Supervised Machine Learning"

### Supplementary Tables and Figures

| Condition | Parameter values |
|---|---|
| Substitution model misspecification | LG-sim+WAG-infer, WAG-sim+LG-infer |
| Recombination rate | Blocks=1,2,3,4 |
| Recombination | Fraction={0.0, 0.1, …, 0.9, 1.0} |
| Lineage-specific heterotachy | Gamma(4, .25) |
| Gene-specific heterotachy | Gamma(4, .25) |

Supplementary Table 1: Additional conditions for simulated test datasets

| **Manuscript** | **Matrix** |
|---|---|
| Borowiec et al. 2015 | Borowiec2015_Best108 |
| Borowiec et al. 2015 | Borowiec2015_Total1080 |
| Nosenko et al. 2013 | Nosenko2013_ribo_11057 |
| Ryan et al. 2013 | Ryan2013_est |
| Ryan et al. 2013 | Ryan2013_est_only_choanozoa |
| Ryan et al. 2013 | Ryan2013_est_only_holozoa |
| Whelan et al. 2015 | Whelan2015_D10 |
| Whelan et al. 2015 | Whelan2015_D10_only_choanozoa |
| Whelan et al. 2015 | Whelan2015_D1_only_choanozoa |
| Whelan et al. 2015 | Whelan2015_D1_only_holozoa |
| Whelan et al. 2015 | Whelan2015_D20_only_choanozoa |
| Whelan et al. 2017 | Whelan2017_full |
| Whelan et al. 2017 | Whelan2017_full_only_choanozoa |

Supplementary Table 2: Metazoa data matrices used in this paper.

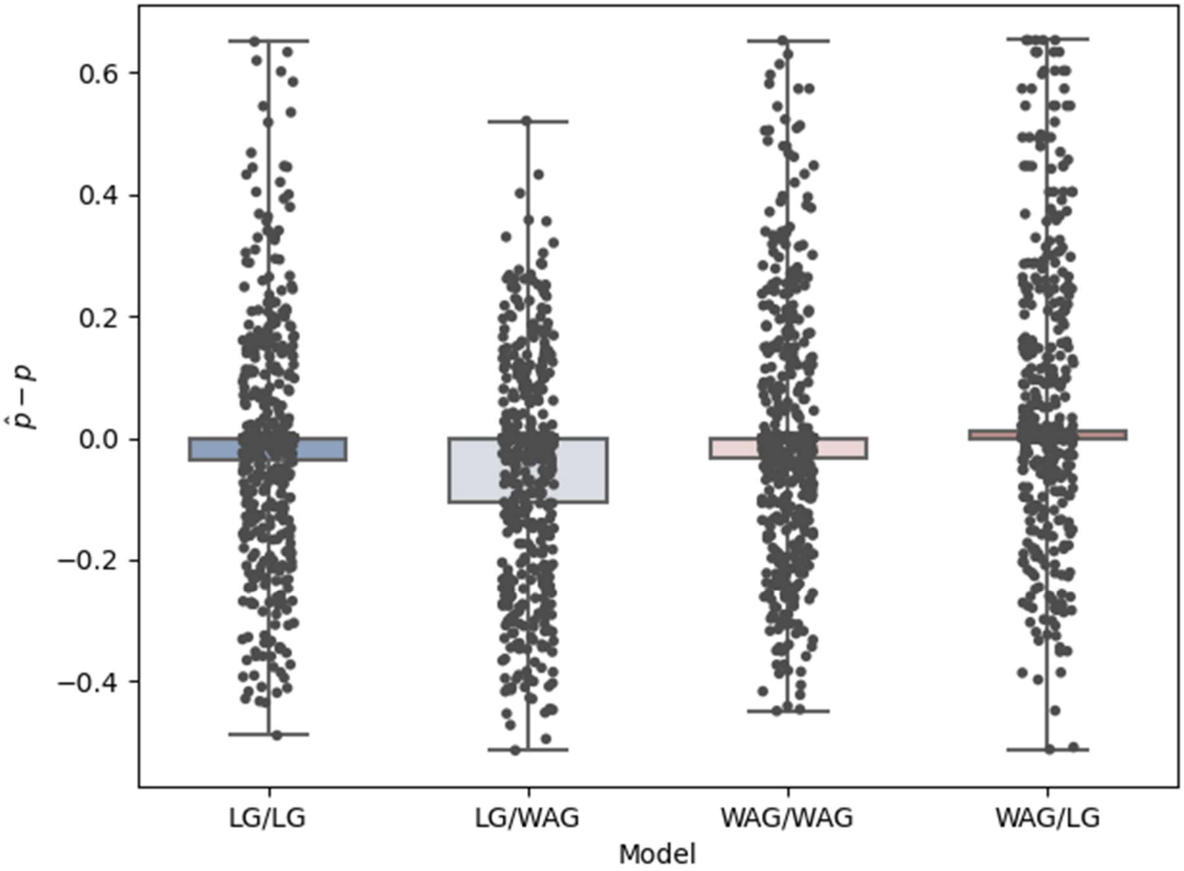


Supplementary Figure 1a: Absolute error in $p$ for mis-specified amino acid substitution matrices. "A/B" labels indicate that alignments were simulated with model "A" and gene trees inferred with substitution matrix "B". Differences in means are not significant (Kruskal-Wallis test, $H = 1.311$, $P = 0.726$).

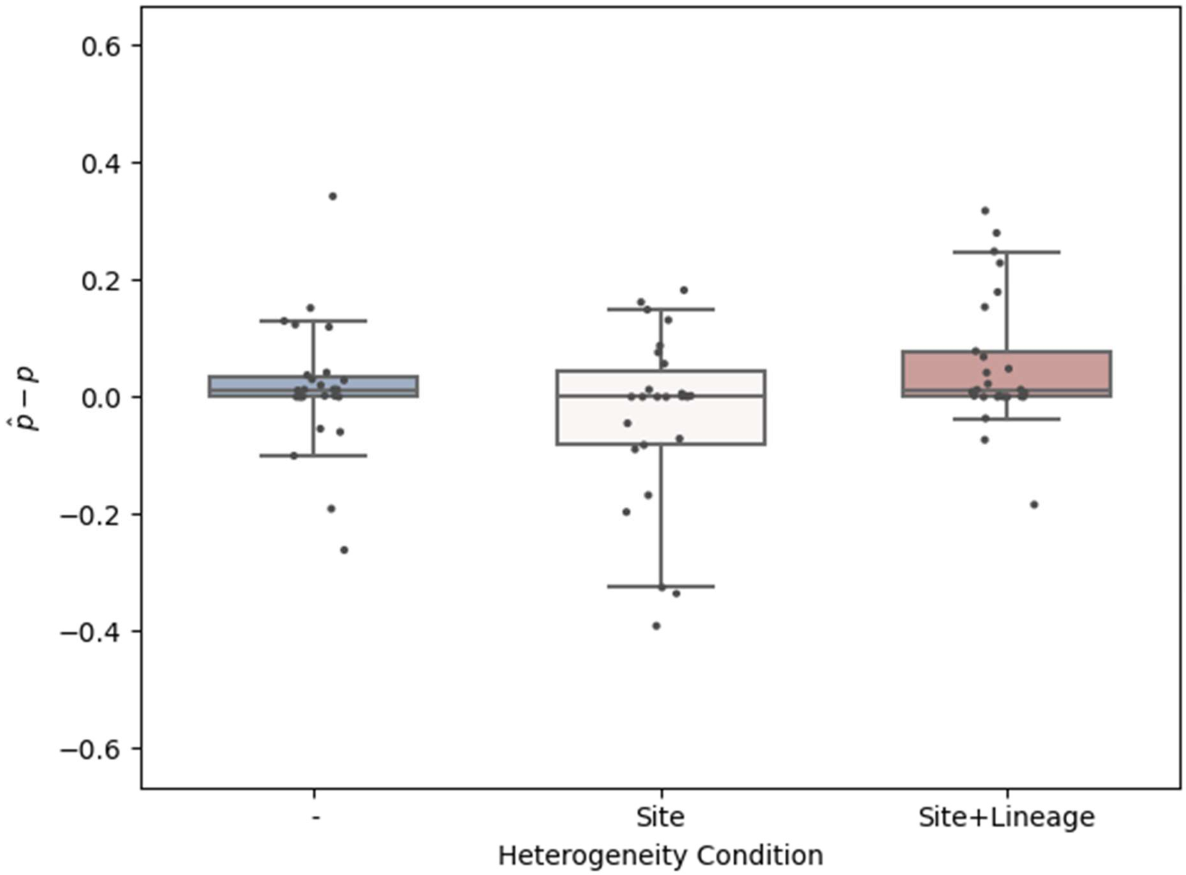


Supplementary Figure 1b: Site-specific rate heterogeneity (middle condition, “Site”) in the test data increases error compared to a single shared rate (left condition, “-”), though this difference is not statistically significant.  Adding lineage-specific heterogeneity (right, “Site+Lineage”) also does not significantly impact error.

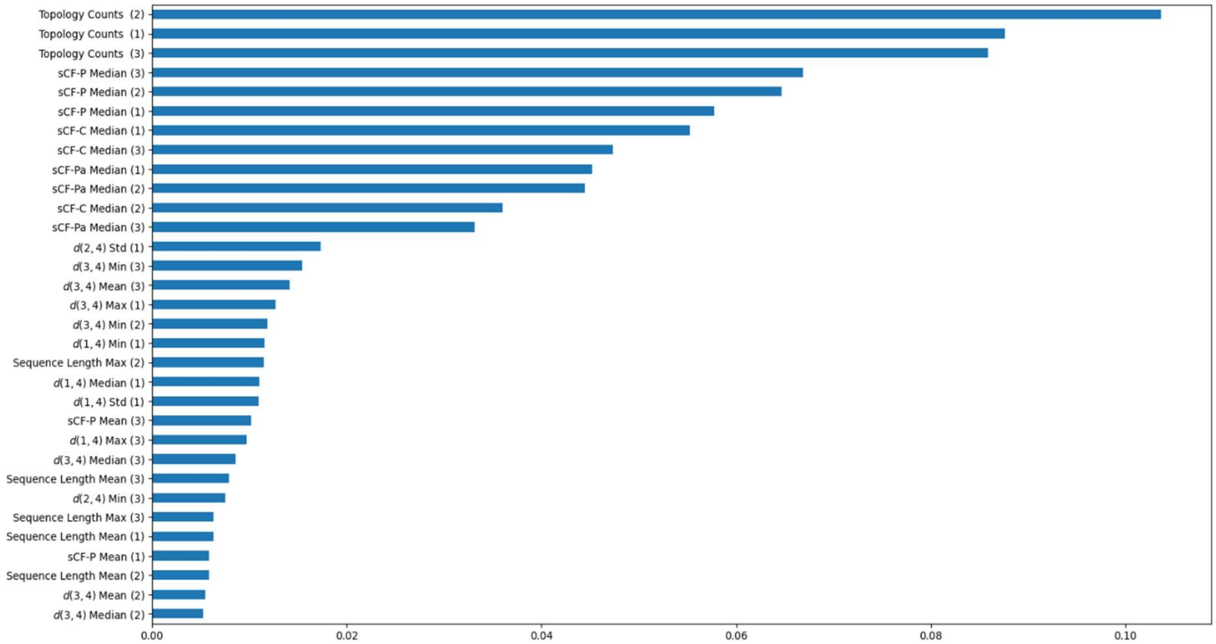
Topology Counts (2)
Topology Counts (1)
Topology Counts (3)
sCF-P Median (3)
sCF-P Median (2)
sCF-P Median (1)
sCF-C Median (1)
sCF-C Median (3)
sCF-Pa Median (1)
sCF-Pa Median (2)
sCF-C Median (2)
sCF-Pa Median (3)
$d(2,4)$ Std (1)
$d(3,4)$ Min (3)
$d(3,4)$ Mean (3)
$d(3,4)$ Max (1)
$d(3,4)$ Min (2)
$d(1,4)$ Min (1)
Sequence Length Max (2)
$d(1,4)$ Median (1)
$d(1,4)$ Std (1)
sCF-P Mean (3)
$d(1,4)$ Max (3)
$d(3,4)$ Median (3)
Sequence Length Mean (3)
$d(2,4)$ Min (3)
Sequence Length Max (3)
Sequence Length Mean (1)
sCF-P Mean (1)
Sequence Length Mean (2)
$d(3,4)$ Mean (2)
$d(3,4)$ Median (2)
0.00
0.02
0.04
0.06
0.08
0.10

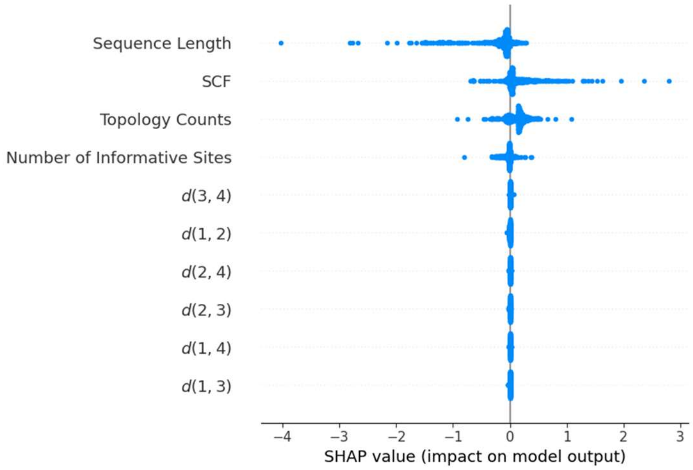
Sequence Length
SCF
Topology Counts
Number of Informative Sites
$d(3,4)$
$d(1,2)$
$d(2,4)$
$d(2,3)$
$d(1,4)$
$d(1,3)$
−4
−3
−2
−1
0
1
2
3
SHAP value (impact on model output)

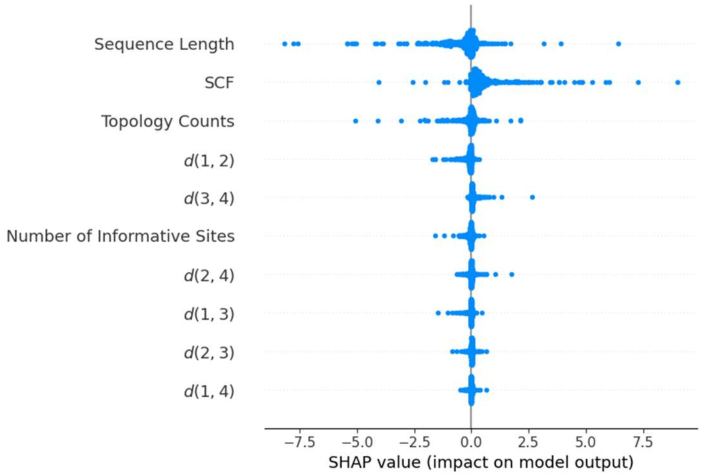
Sequence Length
SCF
Topology Counts
$d(1,2)$
$d(3,4)$
Number of Informative Sites
$d(2,4)$
$d(1,3)$
$d(2,3)$
$d(1,4)$
−7.5
−5.0
−2.5
0.0
2.5
5.0
7.5
SHAP value (impact on model output)

Supplementary Figure 2: a) Gini importance for the top features of the Gradient Boosting *p* regressor. Features are labeled by source (sCF, topology, pairwise distance, etc), summary statistic (Min/Max/ Median/Mean/Std) and topology (1-3). The topology count, being a property of the entire bin, rather than individual gene tree, does not require summary statistics.
b-c) Shapley values for recent (EBL< 0.8) and ancient (EBL>2.0) subsets of the test data for grouped features under DNN-Pred. Features with high dispersion have a higher impact on predictions.

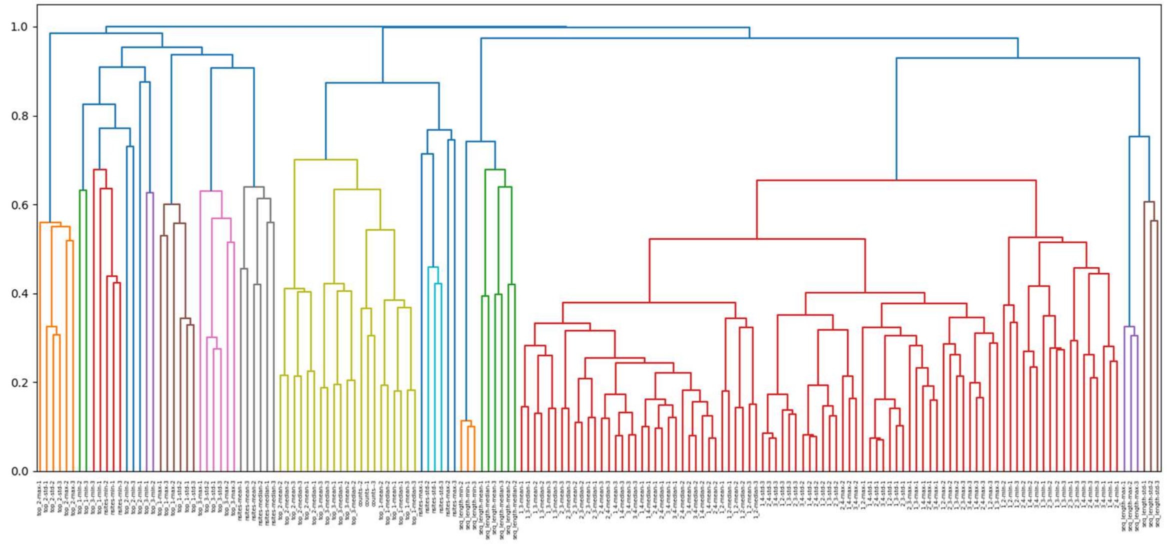


Supplementary Figure 3: *xgboost*-based hierarchical clustering of derived features for the entire training dataset.

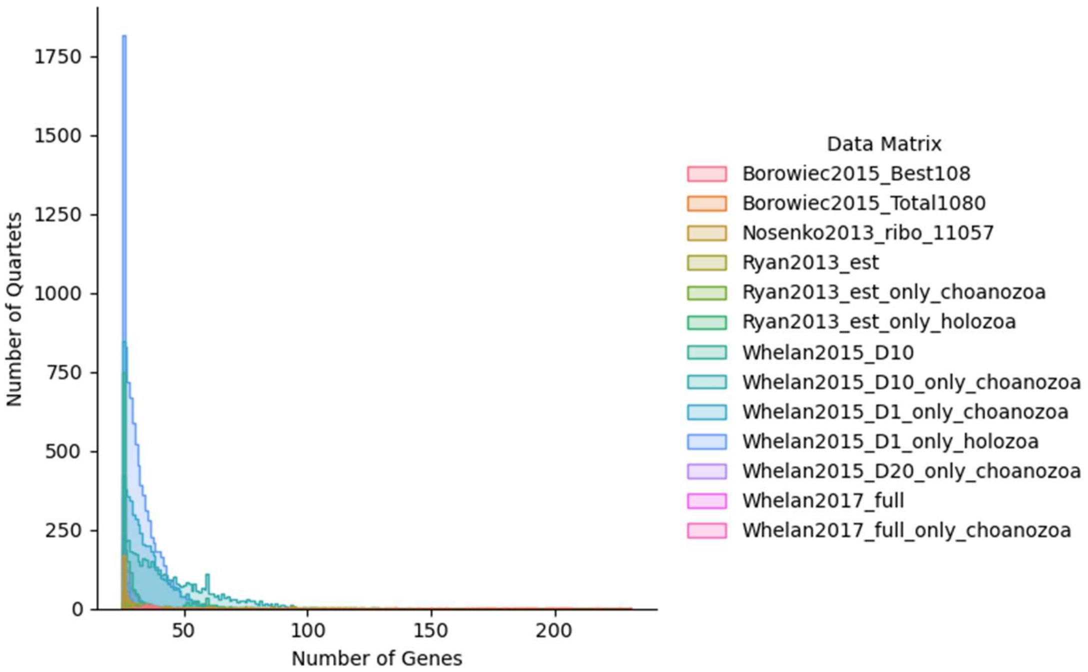


Supplementary Figure 4: Number of genes present for each Ct/Pa/P/Out quartet across all Metazoa datasets.

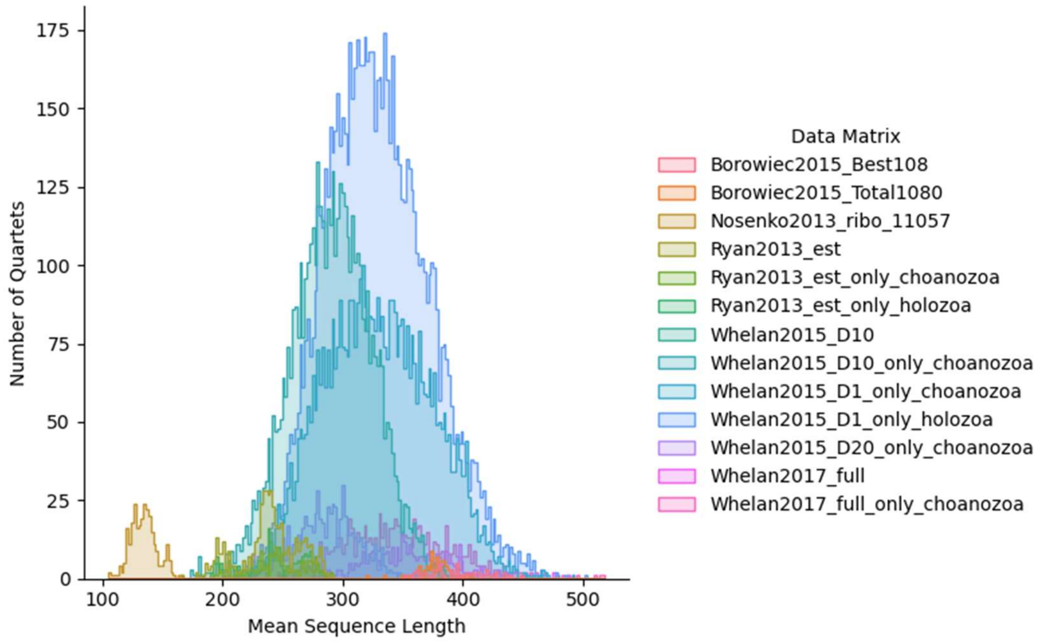


Supplementary Figure 5: Per-quartet sequence length distribution across all Metazoa datasets.

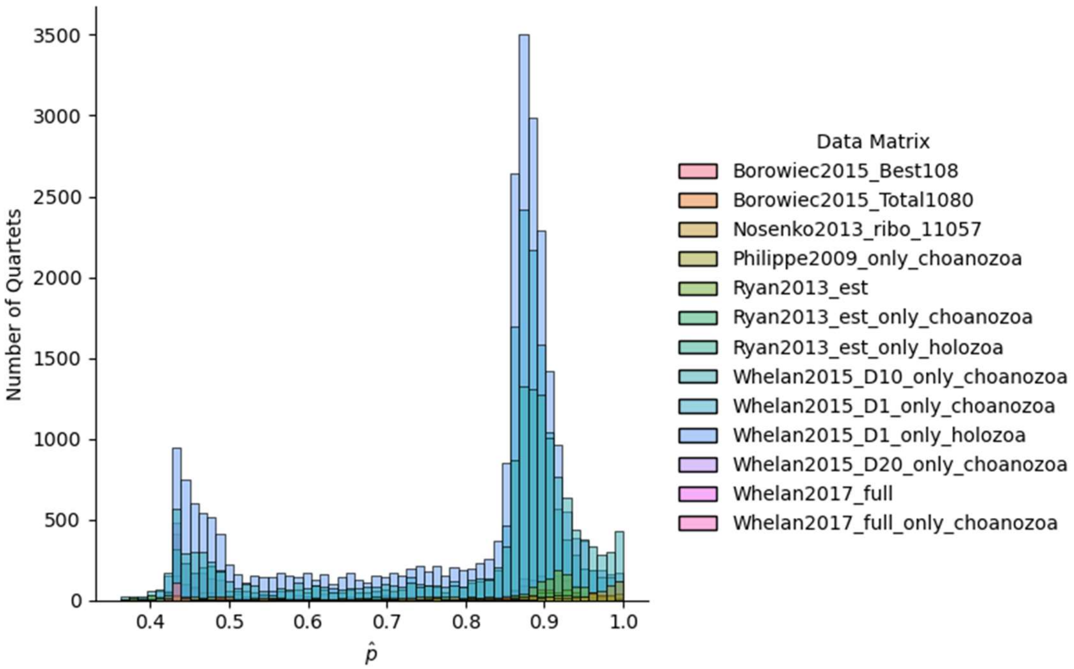


Supplementary Figure 6: DNN-Pred results for 13 Metazoa data matrices. The *y*-axis indicates the number of species quartets within each matrix for which DNN-Pred predicts a particular value of *p*.

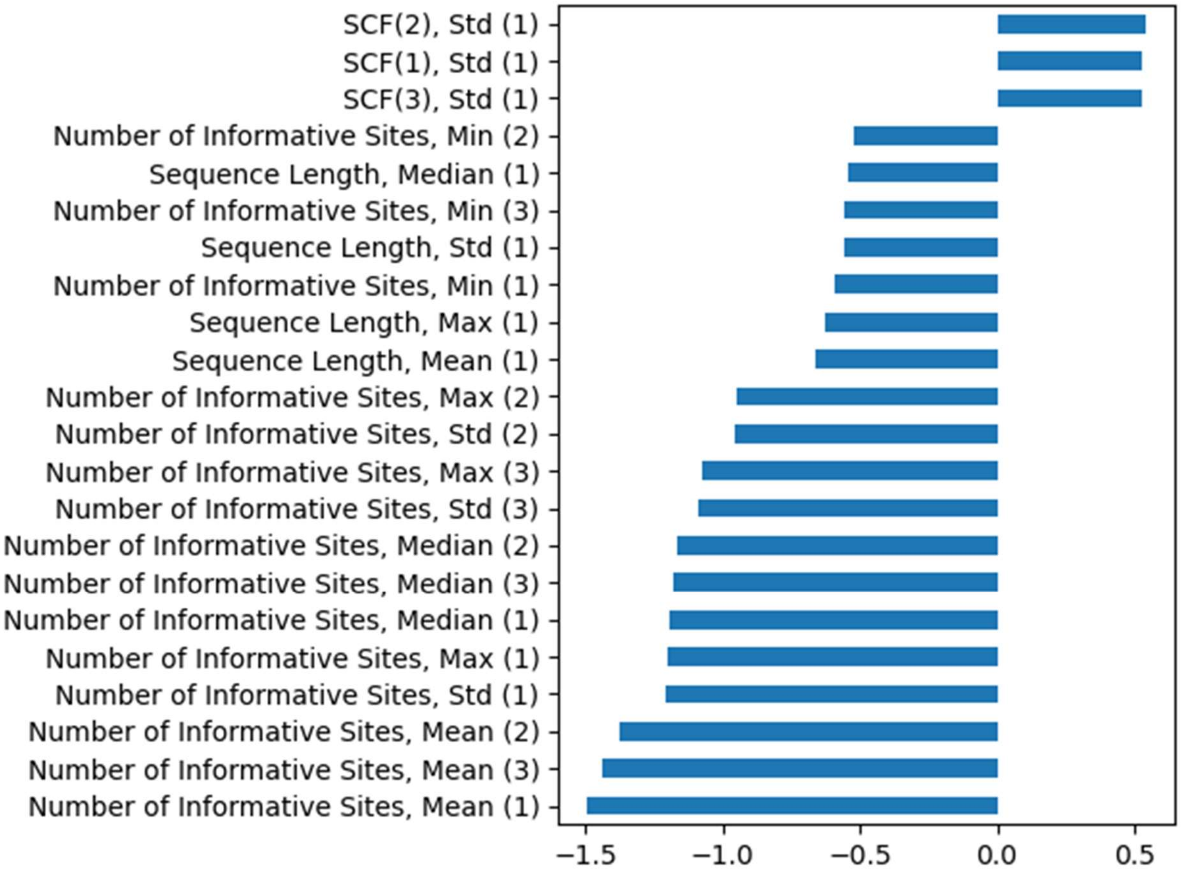


Supplementary Figure 7: Large magnitude Z-scores of feature values for quartets in the 5% and 95% tails of $p$. Positive Z-scores are associated with high predicted $p$, negative Z-scores with low predicted $p$.

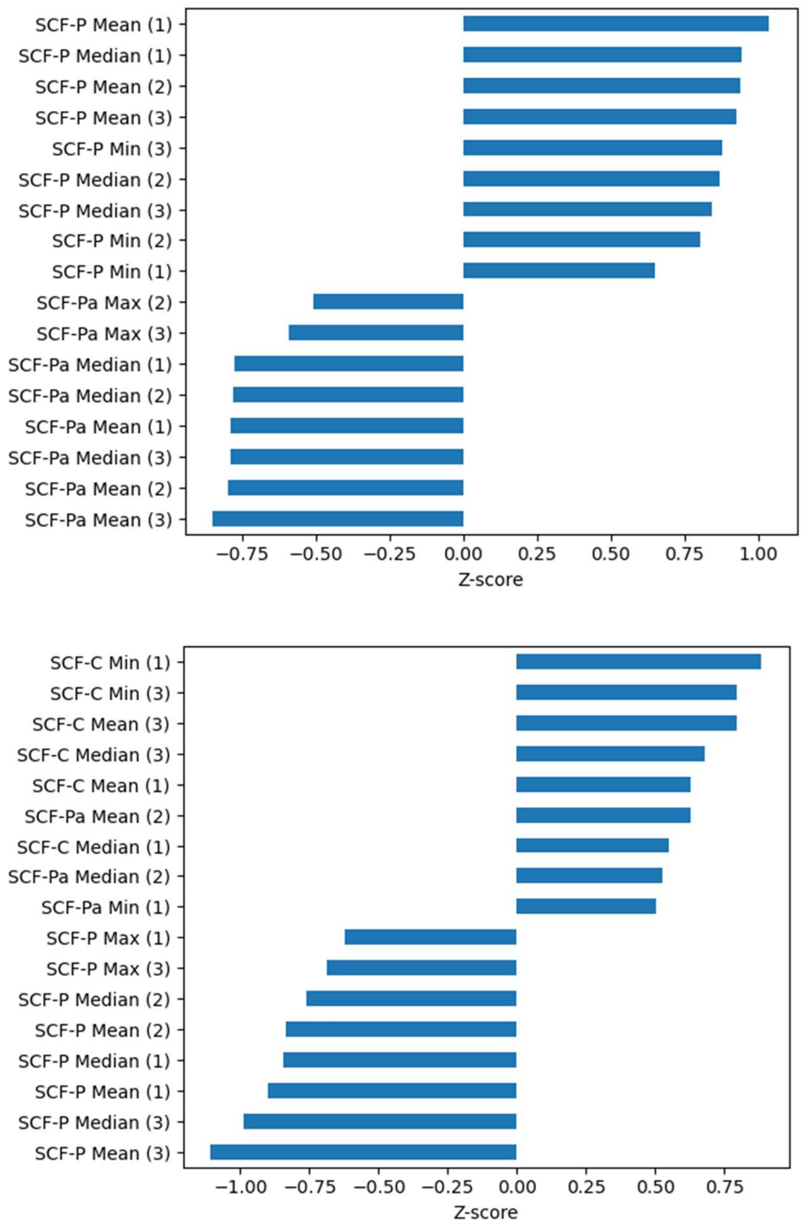


Supplementary Figure 8: Average Z-scores of feature values for all quartets in the 5% and 95% tails of the distribution of likelihood ratio scores for Porifera-sister versus Ctenophora-sister predictions. Features with Z-score values in the interval (-0.5, 0.5) are not shown. a) Quartets which strongly support Porifera-sister have high values of sCF-derived statistics supporting the Porifera-sister topology, and low values of sCF-derived statistics supporting Ctenophore-sister and Parahoxozoa-sister. b) Quartets which support Ctenophore-sister have high values of sCF statistics supporting the Ctenophore-sister topology, and low sCF-derived statistics supporting Porifera-sister.
